# Zelda is dispensable for *Drosophila melanogaster* histone gene regulation

**DOI:** 10.1101/2023.12.19.572383

**Authors:** Tommy O’Haren, Tsutomu Aoki, Leila E. Rieder

## Abstract

To ensure that the embryo can package exponentially increasing amounts of DNA, replication-dependent histones are some of the earliest transcribed genes from the zygotic genome. However, how the histone genes are identified is not known. The pioneer factors Zelda and CLAMP collaborate at a subset of genes to regulate zygotic genome activation in Drosophila melanogaster and target early activated genes to induce transcription. CLAMP also regulates the embryonic histone genes and helps establish the histone locus body, a suite of factors that controls histone mRNA biosynthesis. The relationship between Zelda and CLAMP led us to hypothesize that Zelda helps identify histone genes for early embryonic expression. We found that Zelda targets the histone locus early during embryogenesis, prior to histone gene expression. However, depletion of *zelda* in the early embryo does not affect histone mRNA levels or histone locus body formation. While surprising, these results concur with other investigations into Zelda’s role in the early embryo, suggesting the earliest factors responsible for specifying the zygotic histone genes remain undiscovered.

## Introduction

To efficiently store nearly 2 meters of DNA in a nucleus of only ∼10 micrometers, eukaryotic cells utilize histone-mediated compaction and organization of DNA through the use of nucleosomes, histone protein octamers consisting of two dimers of H2A-H2B and two dimers of H3-H4, a linker histone H1, and the associated ∼150bp of DNA (Annunziato, 2008). The canonical, replication-dependent histones often exist as a multi-gene family with dozens or hundreds of coding sequences (Bongartz & Schloissnig, 2019; Marzluff et al., 2002). The many copies of the histone genes may cluster, but the number of distinct histone loci varies across species with 2 to 4 in most mammals (Seal et al., 2022), 11 in *C. elegans* (Roberts et al., 1987), and 1 in *Drosophila melanogaster* (White et al., 2011). This clustered organization ensures that histone genes are efficiently and quickly expressed to organize the newly synthesized DNA during DNA replication, prior to nuclear/cellular division (Mei et al., 2017; Romeo & Schümperli, 2016). Disruption of proper histone gene expression such that too few or too many histones are produced can have disastrous effects: in the rapidly-dividing developing metazoan embryo, where histones are especially important, a reduction of histones slows cell divisions, whereas histone overexpression causes aberrant divisions and perturbs cell cycle timing. Both conditions lead to embryonic lethality (Amodeo et al., 2015; Chari et al., 2019).

Metazoan embryogenesis represents a period of rapid nuclear/cellular divisions, as quickly as every 8 minutes in the fruit fly (McCleland, Shermoen, & O’Farrell, 2009). Initially, to accommodate the doubling of DNA every cycle, the egg is maternally loaded with vast amounts of histone mRNA that are translationally upregulated after fertilization (Horard & Loppin, 2015; Li et al., 2012). This maternal deposition allows the zygote to proceed through development without the burden of producing its own histone transcripts. Eventually, however, when the maternal transcripts are exhausted or degraded, the zygotic genome must take over transcriptional responsibility (Tadros & Lipshitz, 2009). This process of an embryo beginning to transcribe its own genome is called Zygotic Genome Activation (ZGA) and is not specific to the histone genes, but targets early developmental genes (Schulz & Harrison, 2019).

The minor, early wave of ZGA in *D. melanogaster* begins at nuclear cycle 8 and concludes at cycle 12 when the major, later wave begins and the remainder of the genome is activated (De Renzis et al., 2007; Tadros & Lipshitz, 2009). Zygotic histone gene expression is first detectable around nuclear cycle 11 (White et al., 2007). Zygotic histone expression is accompanied by the formation of a phase-separated, nuclear body consisting of a suite of factors responsible for controlling histone mRNA biosynthesis and mRNA processing called the Histone Locus Body (HLB) (Liu et al., 2006; Tatomer et al., 2016). Although some members of the HLB are known and a general understanding of its basic structure exists (Kemp et al., 2021; Salzler et al., 2013), it is unclear how the histone genes are targeted so early during embryogenesis for their unique regulation, as no HLB specific factors interact with DNA sequence.

The *Drosophila* pioneer factor CLAMP binds genome wide (Larschan et al., 2012), but is also important for HLB formation and proper embryonic expression of the histone genes (Rieder et al., 2017). CLAMP is currently the only known *Drosophila* HLB member, aside from general transcription factors, with DNA binding capability (Tatomer et al., 2016): it recognizes GA-repeat *cis* elements present in the *histone3/histone4* promoter (Koreski et al., 2020; Rieder et al., 2017). Elsewhere, CLAMP functions with another pioneer factor, Zelda, to control ZGA across the fly genome (Duan et al., 2021). Zelda is considered the “master regulator” of *Drosophila* ZGA, binding early, immediately prior to the minor wave of ZGA (Hamm & Harrison, 2018; Harrison et al., 2011). Zelda’s role at the histone locus is unknown, but prior research revealed an intriguing connection: maternal germline *zelda* RNAi resulted in larger *histone3* RNA FISH and HLB factor puncta in the early embryo (Huang et al., 2021). The above observations led us to hypothesize that Zelda helps to specify the histone genes prior to widespread ZGA, plays a role in HLB formation, and regulates zygotic histone biogenesis.

Here, we demonstrate that Zelda targets several sites in the histone gene array prior to zygotic histone gene expression. However, modulating Zelda’s presence at the histone locus has little effect on histone mRNA levels in the early embryo. *Zelda* depletion by RNAi does not cause detectable changes in HLB formation or histone mRNA levels. Additionally, elimination of Zelda’s DNA binding sites within a transgenic histone gene array does not prevent HLB factor recruitment to the transgene. Overall, we conclude that Zelda has a dispensable role in HLB formation and histone gene regulation in the early *Drosophila* embryo. This finding is surprising given Zelda’s status as the master regulator of ZGA and collaboration with CLAMP. The mechanism for specific, early activation of the histone genes remains undiscovered.

## Results and Discussion

### Zelda localizes to TAGteam sites in the histone gene array early during embryogenesis

Zelda recognizes and binds DNA via a series of C2H2 zinc fingers near its C-terminus(McDaniel et al., 2019). Zelda targets TAGteam sites, which include the seven base pair motif: CAGGTAG(Rahul Satija & Robert K. Bradley, 2012). We identified several likely TAGteam sites within the ∼5 Kb *D. melanogaster* histone gene array. The sequences are present in the promoters and gene bodies of the canonical histone genes **(Fig. 1A)**.

**Figure 1.**
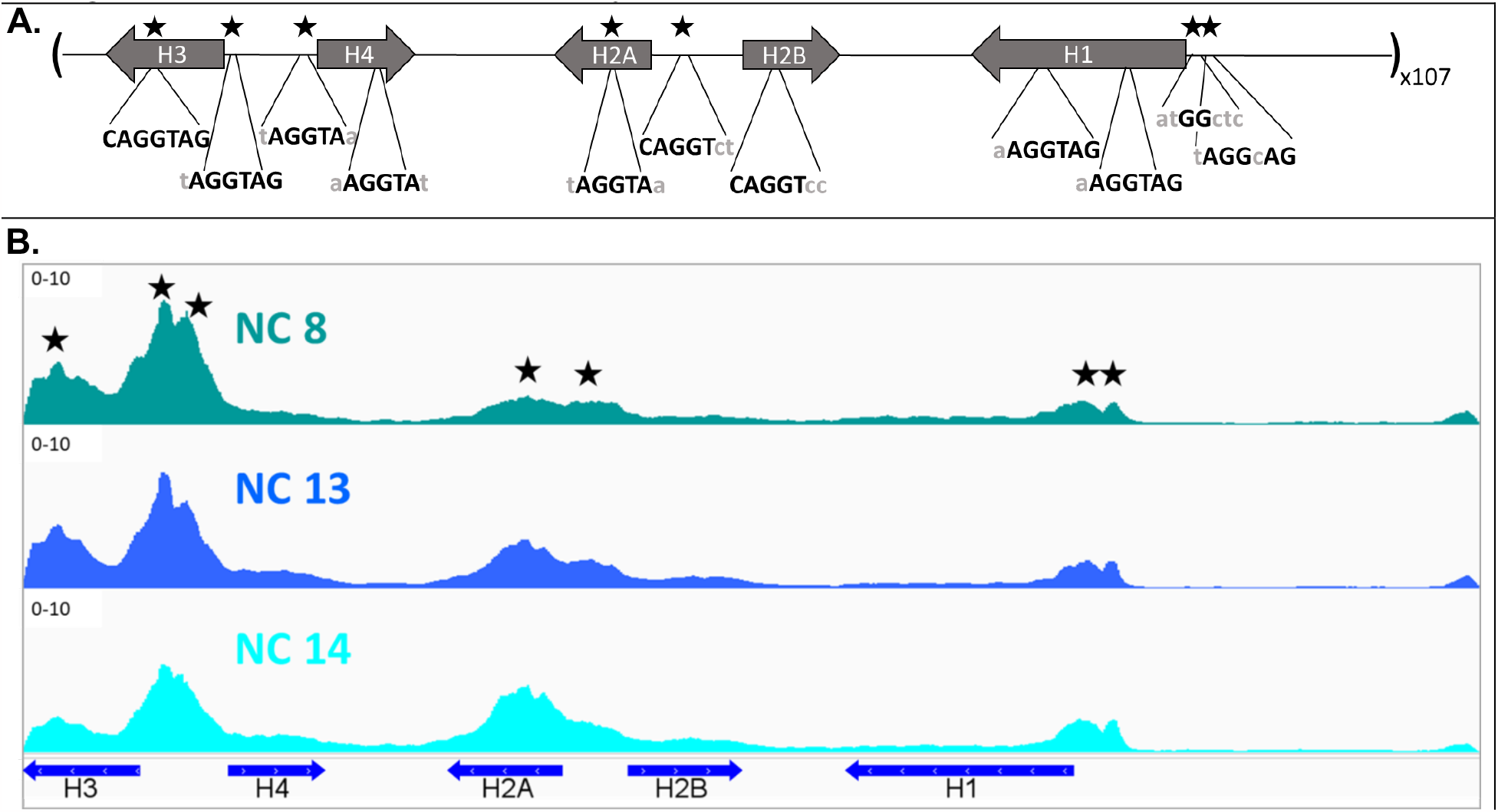
Zelda targets TAGteam sites in the *Drosophila* early embryonic histone gene array. **(A)** The *Drosophila melanogaster* histone locus contains ∼100 tandem repeats of a 5-kb array containing the five canonical replication-dependent histone genes. Derivatives of the TAGteam sequence recognized by the Zelda protein, CAGGTAG, are present across the locus. Seven of these sites correspond to Zelda ChIP-seq signal peaks (Harrison et al., 2011) and are denoted by stars. **(B)** Zelda ChIP-seq shows that Zelda targets the histone gene array as early as nuclear cycle (NC) 8 (teal tracks), immediately after Zelda is translationally upregulated and the start of the minor wave of ZGA. Zelda remains at the array through ZGA, though its presence shifts from predominately at the *histone3/histone4* promoters to the evenly distributed at the former and the *histone2a* gene body (dark blue and light blue tracks).

We began by mapping previously published Zelda ChIP-seq data from early, staged *Drosophila* embryos(Harrison et al., 2011) to the histone gene array. Because the ∼100 histone gene arrays are nearly identical in sequence(Bongartz & Schloissnig, 2019), we can collapse all ChIP-seq data onto a single histone array(McKay et al., 2015). We discovered that Zelda targets the histone gene array by embryonic nuclear cycle 8 **(Fig. 1B)**, the beginning of the minor wave of ZGA and immediately after Zelda is translationally upregulated(McDaniel et al., 2019). Zelda recognizes 7 sites in the array, which correspond to predicted TAGteam motifs **(Fig. 1A)** and targets the histone gene array prior to histone gene expression and HLB factor localization, which is first detectable around nuclear cycle 11(Edgar & Schubiger, 1986; White et al., 2011). Initially, Zelda ChIP signal is highest over sites in the *histone3/histone4* promoter. This promoter is the minimal sequence required for HLB formation and contains important *cis*-regulatory elements targeted by CLAMP(Koreski et al., 2020; Rieder et al., 2017; Salzler et al., 2013). Zelda’s relative distribution across the array changes as development proceeds, though the significance of this distribution, if any, is unclear.

### Zelda reduction in the embryo has little effect on histone transcript levels and HLB factor recruitment

To investigate Zelda’s role in histone gene regulation in the early embryo, we began with an RNAi-mediated depletion (Ni et al., 2011)of maternally deposited *zelda* (Yamada et al., 2019)and measured the effect on histone transcript levels. Using the “maternal triple” GAL4 driver (MTD), we expressed *zelda* shRNA in adult female ovaries and depleted maternally deposited *zelda* mRNA in the egg and early embryo. As previously observed, maternal germline *zelda* RNAi led to nearly 100% embryonic lethality (Duan et al., 2021). Premature Zelda translation does not result in premature gene activation, but alters the levels of transcripts at the normal time of activation (Larson et al., 2022).

When quantified through qPCR, we found that *zelda* RNAi led to significantly reduced levels of *zelda* in 2-4 hour embryos (2-4 hour post-lay), less than 10% of control *mCherry* RNAi levels **(Fig. 2A)**. However, *zelda* depletion did not result in meaningful changes in histone mRNA levels in post-ZGA embryos. Although, we did observe a slight increase in *histone4* mRNA levels. Our observations regarding *zelda* RNAi are very different from our published observations regarding *clamp* RNAi (Rieder et al., 2017), which gives a similarly striking reduction of *clamp* in the early embryo but also leads to significantly decreased levels of multiple histone mRNA levels in post ZGA embryos **(Fig. 2A)**. We also observe the documented upregulation of *zelda/clamp* levels in the embryo when the reciprocal factor is depleted (Duan et al., 2021), serving as additional confirmation of RNAi efficacy.

**Figure 2.**
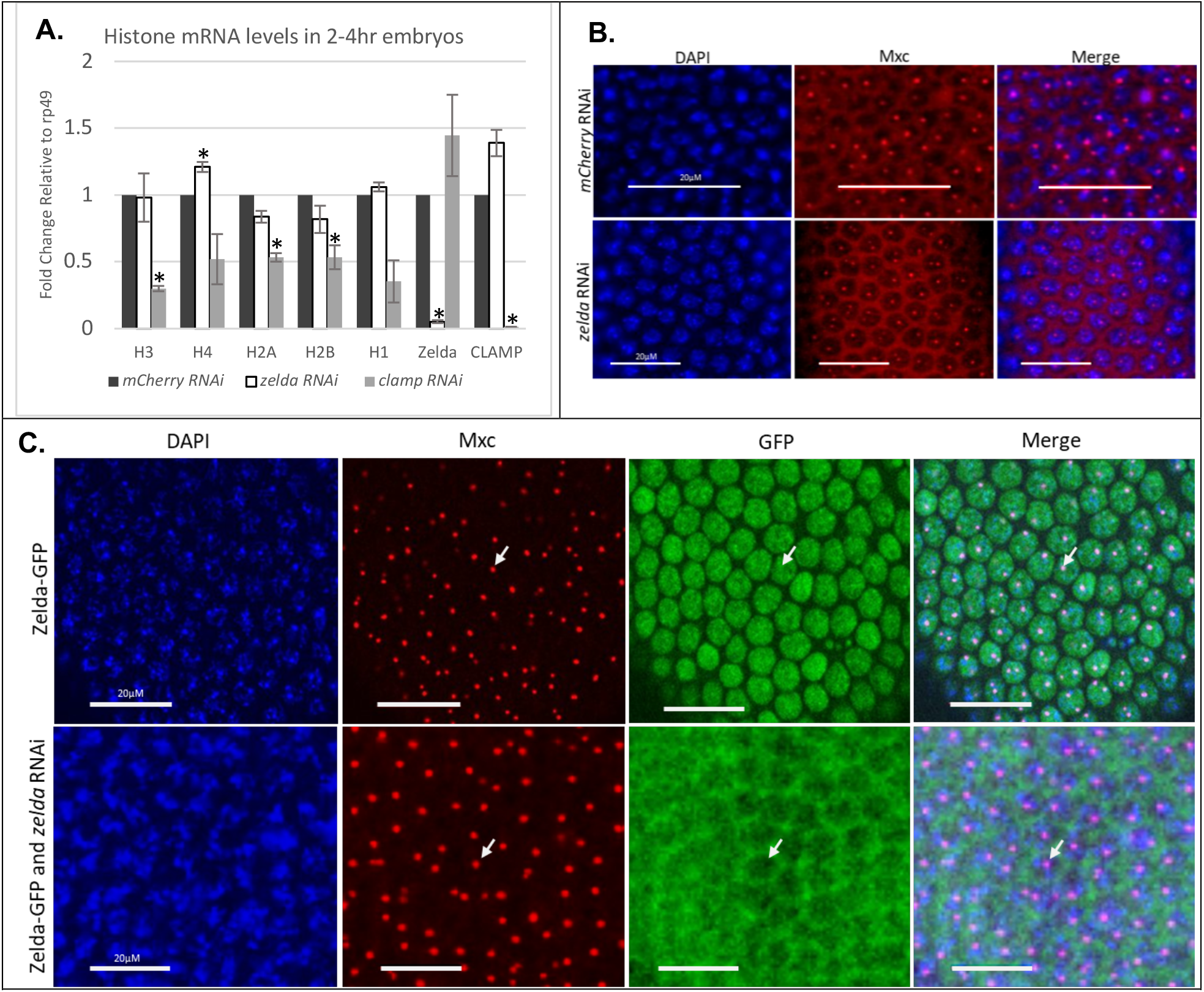
Depletion of *zelda* in the early embryo does not significantly affect histone mRNA levels or HLB factor recruitment. (A) We performed qRT-PCR for histone and transcription factor mRNA in 2-4 hour post lay (after the start of ZGA and zygotic histone gene expression) embryos under control (*mCherry* RNAi), Zelda depleted (*zelda* RNAi), and CLAMP depleted (*clamp* RNAi) conditions. We observe little to no change in histone mRNA levels in the *zelda* depleted embryos compared to controls outside of a slight increase in *histone4* mRNA levels (* denotes a p-value≤0.05, Student’s t-test). In contrast, we see significantly decreased histone transcripts in our *clamp* depleted conditions compared to controls (* denotes a p-value≤0.05, Student’s t-test). Error bars represent ± standard error. Expression is normalized to *rp49*. (B) We performed immunostaining for the HLB component, Mxc, in early embryos in control (*mCherry)* and zelda depleted conditions. We observed no difference in Mxc. **(C)** We utilized a GFP-tagged Zelda observe potential localization of Zelda in embryos. We did not see clear colocalization between Mxc and Zelda in control (*mCherry)* or *zelda* depleted conditions, in contrast to previous results with CLAMP (Rieder et al., 2017).

Although *zelda* depletion does not greatly affect histone transcript levels, we reasoned that it might affect HLB formation. We therefore performed early embryo immunostaining for Mxc, a core scaffolding HLB factor that is one of the first present at the locus (Terzo et al., 2015; White et al., 2011), to detect body formation under *zelda* depletion conditions. The HLB is visible as either 1 or 2 Mxc puncta in wild-type zygotic nuclei, depending on whether homologous chromosomes are paired or unpaired **(Fig. 2B)**. *Zelda* RNAi embryos die around ZGA but progress far enough through development that we can perform immunostaining to detect HLB formation. We observe no change in HLB puncta formation with regards to presence and qualitative puncta size in *zelda* RNAi embryos, compared to control *mCherry* RNAi embryos **(Fig. 2B)**. Our observations contrast with previously published observations that *histone3* RNA FISH and Mxc puncta increase in diameter upon *zelda* RNAi (Huang et al., 2021).

Zelda therefore appears to be dispensable for zygotic histone gene expression and HLB formation in the embryo, despite targeting the histone gene array prior to HLB formation. We were surprised by these conclusions, given that Zelda regulates key embryonic genes during ZGA. However, Zelda is implicated in multiple regulatory pathways in the early embryo, including the establishment of three-dimensional genome organization (Hug et al., 2017). Loss of Zelda during development leads to loss of insulation in a locus-specific manner. It is possible that Zelda has a role in organizing the histone locus and does not directly affect transcription of the histone genes or recruitment of HLB factors. This scenario could explain the published observations by Huang *et al*., who observed increased Mxc puncta size after *zelda* depletion. However, we did not replicate this phenomenon in our RNAi experiments. Conversely, it may be more likely that the continued presence and increased size of HLBs in *zelda* depletion embryos are because other Zelda-dependent regions of the genome remain inactivated. Additional support for this model comes from a recent study that demonstrates that *zelda* depletion affects RNA Polymerase II clustering across the nucleus, except at the replication-dependent histone genes (Cho et al., 2022).

### Zelda and CLAMP do not affect reciprocal localization to the histone genes

We previously demonstrated that CLAMP targets the *histone3/histone4* promoter by embryonic nuclear cycle 10 (Rieder et al., 2017). Zelda and CLAMP bind targets across the genome during ZGA and regulate each other’s binding at a subset of promoters genome-wide. The binding motifs of Zelda and CLAMP (TAGteam sites and GA-repeats, respectively) are often found in close proximity, within 2kb, across the genome (Duan et al., 2021). We previously performed *zelda* and *clamp* RNAi in early embryos and reciprocal ChIP-seq for the other factor. We mapped these early embryo ChIP-seq datasets to the histone gene array to discover if these two pioneer factors target the locus in collaboration. Upon *zelda* RNAi, CLAMP localization to the histone array remains relatively unchanged with a strong peak at the *histone3/histone4* promoter. Under *clamp* RNAi conditions, CLAMP binding at the locus decreases and changes its distribution across the locus. Under *zelda* RNAi conditions, Zelda is largely lost at the genes, although some small amount appears to remain. Upon *clamp* RNAi, Zelda’s localization to the locus remains mostly unchanged, however, there is a slight drop in binding near the *histone1* gene **(Fig. S1)**. Overall, the loss of one factor does not greatly affect the binding of the other at the histone genes, which conflicts with the relationship elsewhere in the genome where the two play a reciprocal role in the other’s localization and activity(Duan et al., 2021).

We previously documented that even though maternal *clamp* RNAi reduced CLAMP to undetectable levels in the early embryo by western blot, small amounts of maternal CLAMP are still visible, specifically at the histone locus, by immunostaining in the early embryo (Rieder et al., 2017). We therefore hypothesized that a small amount of Zelda might also be detectable at the zygotic histone locus by immunofluorescence. We performed post-ZGA embryo immunofluorescence for Zelda using an endogenously CRISPR-tagged Zelda-GFP (D. C. Hamm et al., 2017) under *mCherry* and *zelda* RNAi conditions. Although we confirmed effective *zelda* knock-down by qPCR **(Fig. 2A)**, we did not observe enrichment of Zelda-GFP signal specifically overlapping with Mxc in either condition; Zelda is broadly nuclear under control (*mCherry* RNAi) conditions, and the majority of this nuclear signal is lost after *zelda* depletion **(Fig. 2C)**. Although we do not detect Zelda at the histone locus by immunofluorescence, we did observe traces of Zelda through ChIP-seq after *zelda* RNAi **(Fig. S1)**, indicating that small amounts of Zelda may remain at the locus even after depletion.

Regulation of the histone genes and the production of sufficient histone proteins is an exceptionally important process, especially in a rapidly dividing/replicating embryo, so nuclei may implement strategies to increase localization and retention of imperative factors. Our findings may explain why *zelda* depletion has little effect on histone transcript levels, as a small amount of Zelda that escapes RNAi may still localize to the histone genes and be sufficient for inducing gene expression. However, we do not observe Zelda colocalizing with Mxc even under control conditions, in contrast to CLAMP, even though small amounts of Zelda are detectable at the histone genes via ChIP-seq. Zelda and CLAMP may have redundant activity or Zelda may have extraneous activity at the histone locus. From our results we further conclude that Zelda likely has dispensable activity at the zygotic histone locus.

### Transgenic histone gene arrays lacking TAGteam sites still recruit HLB factors

Depleting *zelda* in the embryo has broad genome-wide consequences and results in near 100% embryonic lethality. To isolate the effect of Zelda specifically at the histone genes, without pleiotropic effects, we manipulated Zelda’s target elements in a transgenic histone gene array. The relationship between Zelda and its DNA binding motif, the TAGteam sites, is well documented (Foo et al., 2014; Liang et al., 2008; Nien et al., 2011; R. Satija & R. K. Bradley, 2012). The number of TAGteam sites within a region is directly proportional to the response to Zelda: more sites lead to more robust Zelda-dependent activation (Dufourt et al., 2018). Additionally, slight differences in the binding site sequence influence Zelda affinity and activity. Removal of TAGteam sites in certain genes directly eliminates activation of Zelda targets during early development (Li & Eisen, 2018). To eliminate the activity of Zelda specifically at the histone genes, we manipulated the TAGteam sites. It would be extremely difficult, nigh impossible, to edit the endogenous histone locus and the over 100 nearly-identical histone gene arrays (Bongartz & Schloissnig, 2019). Instead, we leveraged a transgenic system in which we can manipulate the sequence of histone array transgene outside of the larger locus to determine sequence features of the array that are required for histone gene regulation. The wild-type version of this transgene recruits all known HLB factors and expresses histone genes similar to the endogenous locus (Koreski et al., 2020; McKay et al., 2015; Rieder et al., 2017; Salzler et al., 2013).

We generated a transgenic histone gene array in which we mutated 4 of the highest bound TAGteam sites to eliminate Zelda binding **(Fig. 3A)**. We were unable to manipulate two of the TAGteam sites due to their position in the array, including a site in the *histone4* promoter that overlaps with the TATA box, as these mutations could affect transcription which is necessary for full HLB formation (Salzler et al., 2013). We chose to eliminate the GG dinucleotide present in the motif because this is the most conserved and most consequential positions(Liang et al., 2008). We inserted the transgene into two genomic locations, one on Chromosome 3L (VK33) and one on Chromosome 3R (Zh86-Fb), using PhiC31 integrase, as chromatin context affects HLB formation at transgenic arrays (Salzler et al., 2013). As positive controls, we used lines carrying wild-type histone array transgenes at the same genomic sites.

**Figure 3.**
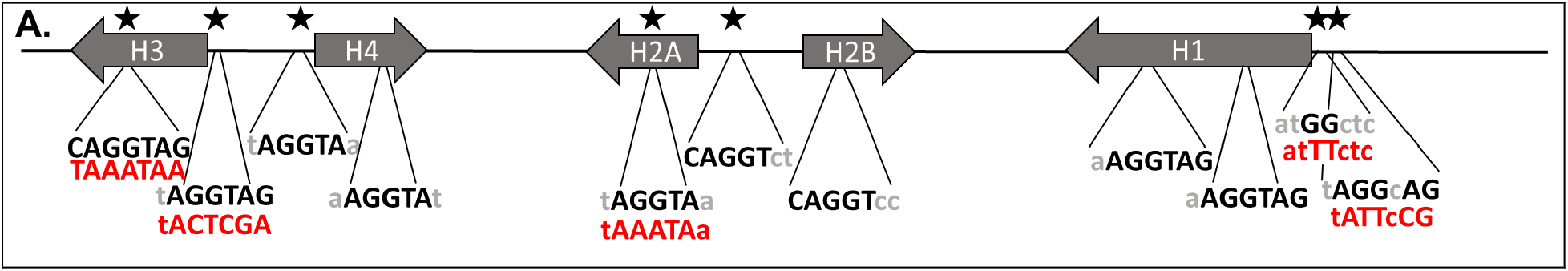
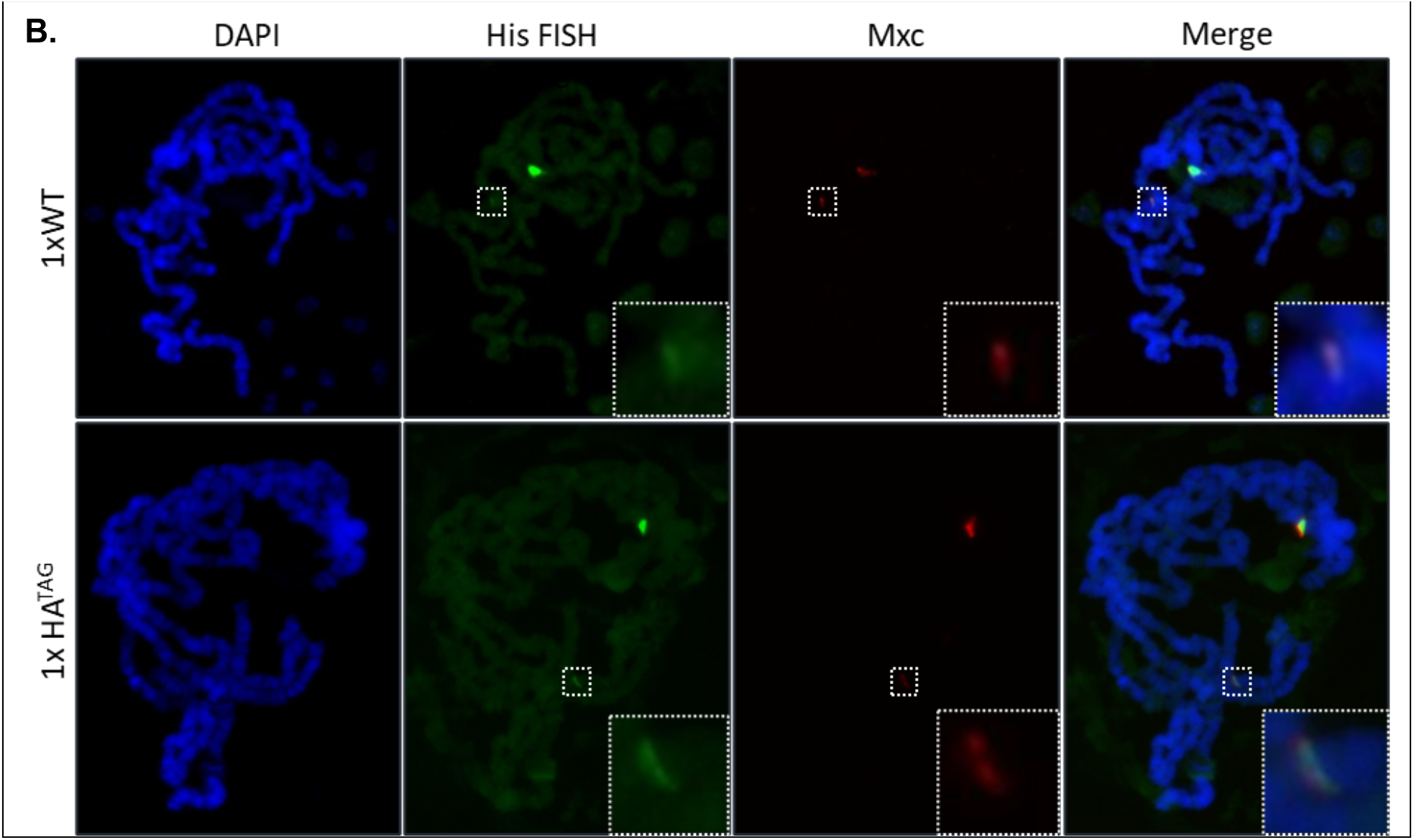
Decreasing Zelda binding sites in transgenic histone arrays does not affect HLB factor recruitment. **(A)** We created a single histone array transgene which is not recognized by the Zelda protein (1xHA^TAG^). Red letters represent the changes to the endogenous sequence after mutagenesis. **(B)** We performed polytene chromosome immunostaining on chromosomes extracted from salivary glands of flies homozygous for a single copy transgenic histone array that matches the endogenous locus sequence (1xWT) or the 1xHA^TAG^ array. We stained for the unique HLB factor Mxc and also performed DNA FISH using probes targeting histone gene sequence to identify the endogenous loci and confirm our transgenic arrays. We observe that an ectopic Mxc fluorescent band is visualizable for both the 1xWT transgene and the 1xHA^TAG^ transgene indicating that the loss of these specific Zelda binding sites does not affect HLB factor recruitment.

We tested the ability of the transgenes to recruit HLB factors by performing polytene chromosome immunostaining, as previously documented (Koreski et al., 2020; Rieder et al., 2017; Salzler et al., 2013). We visualized HLB factor recruitment through staining for the HLB-specific scaffolding protein Mxc (Kemp et al., 2021; Terzo et al., 2015). Recruitment to a single copy of the histone gene array cannot be visualized in embryos. To circumvent this issue, we use polytene chromosome immunostaining as a read out of successful HLB establishment in the embryo. Although Zelda is not expressed in larval salivary glands, the HLB is established in the early embryo, and Mxc remains associated with chromatin through cell cycles/divisions (White et al., 2011).

Our observations demonstrate that Mxc is recruited to the transgenic arrays lacking TAGteam sites (HA^TAG^), similar to the wild-type transgenic arrays (HA^WT^). A single, large band near the chromocenter representing the endogenous histone locus, is visible on all polytene chromosome spreads, and serves as an internal staining control. We also observe a much smaller band representing the single array transgenes, both HA^WT^ and HA^TAG^, at the locations we targeted using the PhiC31 integrase **(Fig. 3B)**. We obtained similar results when we stained for other members of the HLB, Mute and FLASH **(Fig. S2)**. Mute is a negative regulator of histone mRNA expression (Bulchand et al., 2010), while FLASH is a core HLB component involved in transcript 3’ end processing (Duronio & Marzluff, 2017). As previously observed, the integration location of the transgene affects factor recruitment (Günesdogan, Jäckle, & Herzig, 2010; Salzler et al., 2013). We noticed a slight difference in fluorescence in the two transgenic locations: both the HA^WT^ and HA^TAG^ transgenes have decreased fluorescence at VK33 compared to ZH86-Fb. However, we found no detectable difference in the recruitment of HLB factors to HA^TAG^ transgenes compared to HA^WT^ transgenes at the same insertion site **(Fig. S3)**.

The *histone3/histone4* promoter sequence is targeted by both CLAMP (Rieder et al., 2017) and Zelda **(Fig.1)** in the early embryo. To confirm our earlier observation the *zelda* depletion does not impact CLAMP targeting to this region, we performed an electrophoretic mobility shit assay (EMSA). CLAMP is a member of the Late Boundary Complex, which forms during embryogenesis and is detectable by EMSA with embryo extract (Kaye et al., 2017). Probes carrying the wild-type *histone3/histone4* region shift dramatically when exposed to embryo extract **(Fig. S4)**, indicating interaction with proteins in the extract. Deleting the GA-repeats, which interact with CLAMP, from the probe abrogate this interaction. However, elimination of both TAGteam sites did not prevent the shift indicating that the embryonic interaction at this region is preserved in the absence of Zelda at the TAGteam sites. We again conclude that Zelda is likely not involved in recruiting factors to the histone genes **(Fig. S4)**.

### Zelda is likely dispensable for histone gene regulation

The 5 kb histone gene array includes at least 6 TAGteam sites that are targeted by Zelda. The TAGteam sites have slightly different sequences **(Fig. 1A)**, which may affect Zelda binding affinity. It is possible that certain TAGteam sites are more important than others within the locus, likely exemplified by the difference in binding at each site by Zelda ChIP-seq and distribution changes across the array as development proceeds. Our transgenic histone array with mutated Zelda binding sites showed no detectable effect on HLB factor recruitment. However, it may be possible that the two TAGteam sites that remained in our array **(Fig. 3A)** still recruit Zelda at sufficient levels to influence histone gene regulation.

It is also possible that Zelda is important at the locus only under certain unique situations. For example, a histone array transgene lacking CLAMP-recruiting GA-repeat sequences is not targeted by HLB factors or expressed unless the endogenous histone locus is missing, indicating a “backup” or secondary mechanism of locus identification (Koreski et al., 2020; Rieder et al., 2017). While CLAMP does not target a specific region of this transgenic histone gene array by ChIP-seq, it is present in the body by immunofluorescence, suggesting that CLAMP is recruited even in the absence of DNA binding. The TAGteam sites and Zelda may be important in this unique context for CLAMP recruitment, HLB formation, and histone expression. The GA-repeats are poorly conserved in other *Drosophila* species, although CLAMP is still recruited to histone loci (Rieder et al., 2017; Xie et al., 2022). The TAGteam sequences may be more critical in the histone gene arrays of other *Drosophila* species compared to *melanogaster*. Therefore, Zelda may also be more critical at the locus in non-*melanogaster* species, either to recruit CLAMP or as an independent factor.

Overall, we conclude that although Zelda and CLAMP collaborate elsewhere in the genome during ZGA, Zelda is dispensable for proper histone gene regulation and histone locus body formation, unlike CLAMP. CLAMP remains the only known sequence-specific binding factor in *Drosophila* that influences histone expression and HLB formation.

## Materials and Methods

### Drosophila strains

We used the maternal-triple-driver (“MTD”) Gal4 stock (Bloomington, #31777) and a stock expressing shRNA against *zelda* from the Rushlow Lab (Sun et al., 2015) and stocks expressing shRNA against *clamp* (Bloomington, #57008) and *mCherry* (Bloomington, #35787). The Zelda-sfGFP stock was gifted from the Harrison Lab (Danielle C. Hamm et al., 2017)and was crossed to the shRNA *zelda* strain to generate the combined line that allowed the knock-down of the tagged protein. We maintained flies on standard cornmeal/molasses food. We maintained GAL4 crosses at 24°C and crosses/stocks for polytene chromosome preparation at 18°C.

### Cloning and transgenesis

To generate the HA^TAG^ histone array transgene, we inserted a wild-type histone array sequence containing the 5 replication-dependent histone genes into a pBluescript II KS+ vector (Agilent #212207) and introduced mutations using a Q5 Site-Directed PCR Mutagenesis Kit (New England Biolabs). After mutagenesis, we transferred the array to the pMulti-BAC vector ((McKay et al., 2015)), sequence confirmed using whole plasmid sequencing (Plasmidsaurus), and inserted into the VK33 (Chr 3L) and Zh86-Fb (Chr 3R) insertion sites using φC31-mediated integration (Genetivision).

### ChIP-seq data analysis and visualization

We analyzed the Zelda ChIP-seq data sets in staged early embryos by directly importing individual FASTQ data sets from Harrison et al., 2011 (NCBI GEO GSE30757) into the web-based platform Galaxy ((Afgan et al., 2016)The Galaxy Community 2023) through the NCBI SRA run selector by selecting the desired runs and utilizing the computing galaxy download feature. We retrieved the FASTQ files from the SRA using the “faster download” Galaxy command. Because the ∼100 histone gene arrays are extremely similar in sequence(Bongartz & Schloissnig, 2019), we can collapse ChIP-seq data onto a single histone array (Hodkinson et al., 2023; McKay et al., 2015; Rieder et al., 2017). We used a custom “genome” that includes a single *Drosophila melanogaster* histone array similar to that in McKay et al. 2015, which we directly uploaded to Galaxy using the “upload data” feature and normalized using the Galaxy command “normalize fasta” specifying an 80 bp line length for the output FASTA. We aligned ChIP reads to the normalized histone gene array using Bowtie2 (Langmead & Salzberg, 2012) to create BAM files using the user built-in index and “very sensitive end-to-end” parameter settings. We converted the BAM files to bigwig files using the “bamCoverage” Galaxy command in which we set the bin size to 1 bp and set the effective genome size to user specified: 5000 bp (approximate size of l histone array). We visualized the Bigwig files using the Integrative Genome Viewer (IGV) (Robinson et al., 2011).

We performed analysis of *zelda* and *clamp* RNAi ChIP-seq data sets from Duan et al., 2021 (NCBI GEO GSE152598) as described above. We used a custom R script to combine and visualize replicates (Xie et al., 2022).

### Quantitative real-time PCR

We performed qRT-PCR as described in (Urban et al., 2017) using RNA extracted from 2-4hr post-lay embryos using Trizol. We performed cDNA synthesis from RNA using LunaScript RT Supermix Kit (New England Biolabs) beginning with 500ng of RNA, which we then diluted 1:20 in MilliQ water prior to PCR. We performed reactions in technical duplicates using the AzuraQuant Green Fast qPCR Mix (Azura Genomics) and the appropriate primers. Primers for each transcript can be found in Table S1. We performed three biological replicates for each genotype and target gene. We performed PCR using the QuantStudio 3 Real-Time PCR System (ThermoFisher Scientific). We normalized transcript abundance to *rp49* and calculated fold change via the ΔΔCt method (Rao et al., 2013). We analyzed data using a Student’s t-test, comparing transcript abundance between *clamp* or *zelda* RNAi embryos to matched *mCherry* control embryos.

### Embryo immunofluorescence

We performed embryo immunofluorescence of fixed, staged embryos after aging embryos laid on grape juice plates to 2-4 hours, which we then dechorionated in 50% bleach and collected using a 40micron strainer. We immediately fixed embryos using 37% formaldehyde in heptane for 20 minutes. We then collected and washed embryos in methanol and stored in -20°C. We began immunostaining by rehydrating embryos using increasing concentrations of PB-Tween in Methanol. We then washed embryos and incubated with primary antibody in block (1% BSA in 1x PBS) overnight at 4°C on a rotator. The following day, we collected embryos and washed in block before incubating with secondary antibody for 2 hours at room temperature protected from light. We then washed and mounted embryos on slides using Prolong Diamond anti fade reagent with DAPI (ThermoFisher Scientific, P36961). We imaged embryos using a Keyence BZ-X810 Fluorescence microscope using a 20x objective. We conducted Image processing using ImageJ software (NIH).

### Polytene chromosome FISH and immunofluorescence

We performed polytene chromosome FISH and immunostaining on chromosomes extracted from salivary glands dissected from third instar *Drosophila* larvae raised at 18°C on standard cornmeal/molasses food. We passed glands through fix 1 (4% formaldehyde, 1% Triton X-100, in 1 × PBS) for 1 min, fix 2 (4% formaldehyde, 50% glacial acetic acid) for 2 min, and 1:2:3 solution (ratio of lactic acid:water:glacial acetic acid) for 5 min prior to squashing and spreading. We washed slides in 1X PBS, then in 1% Triton X-100 (in 1X PBS) and 2 X SSC. We dehydrated slides in ethanol and allowed to air dry. We generated Biotinylated DNA FISH probes for the histone array using a Nick Translation Reaction with biotin-11-dUTP (primers found in Table S1). We then placed the slides on a heating block set to 91°C after applying the biotinylated FISH probe targeting the histone gene array in hybridization buffer (2 X SSC with dextran sulfate, formamide, and salmon sperm DNA) and sealed the coverslip with rubber cement. We incubated slides at 37°C overnight in a humid chamber. We peeled off the rubber cement and washed slides in 2 X SSC to remove coverslips and then washed in 1 X PBS. Next, we blocked for one hour in 0.5% BSA diluted in 1X PBS. We then incubated slides with primary antibodies diluted in blocking solution (antibodies specifics below) overnight at 4°C in a dark, humid chamber. We washed slides in 1 X PBS and 2 X Tween-20/NP-40 wash buffers. We next incubated the slides with streptavidin-488 (DyLight)in detection solution for 1hr in a humid chamber and then washed in 1 x PBS. We incubated slides with secondary antibody diluted in blocking solution (antibody specifics below) for two hours at room temperature. We washed and mounted slides in Prolong Diamond anti fade reagent with DAPI (ThermoFisher Scientific, P36961), and imaged chromosome spreads on a Zeiss Scope.A1 equipped with a Zeiss AxioCam using a 40×/0.75 plan neofluar objective using AxioVision software. We conducted image processing using ImageJ software (NIH).

### Antibodies

We used primary antibodies at the following concentrations: guinea pig anti-Mxc (1:5000; gift from Drs. Robert Duronio and William Marzluff), rabbit anti-GFP (1:1000; ThermoFisher Scientific #A-6455), rabbit anti-FLASH (1:2000, gift from Drs. Robert Duronio and William Marzluff), guinea pig anti-Mute (1:5000; Bulchand et al. 2010). We used secondary antibodies (ThermoFisher Scientific) at a concentration of 1:1000: goat anti-guinea pig AlexFluor 647 (#A-11073), rabbit anti-guinea pig TRITC(#PA1-28594), goat anti-rabbit AlexFluor 488 (#A-21450).

### EMSAs and probes

We performed EMSAs as described in (Aoki et al., 2008) with some modifications. Late embryo nuclear extracts were prepared from 6-18 hour Oregon R embryo collected on apple juice plates and aged 6 hours at room temperature. We performed nuclear extract preparations as in Aoki et al.; however, we omitted the final dialysis step and completed the extraction with the final concentration of KCl at 360 mM. We made EMSA probes using PCR using primers found in Table S1. We 5’ end labeled one pmol of probe with γ-32P-ATP (MP Biomedicals) using T4 polynucleotide kinase (NEB) in a 50µl total reaction volume at 37°C for 1 hour. We used Sephadex G-50 fine gel (Amersham Biosciences) columns to separate free ATP from labeled probes. We adjusted the volume of the eluted sample to 100 μl using deionized water so that the final concentration of the probe was 10 fmol/μl. We performed 20 μl binding reactions consisting of 0.5 μl (5 fmol) of labeled probe in the following buffer: 25 mM Tris-Cl (pH 7.4), 100 mM KCl, 1 mM EDTA, 0.1 mM dithiothreitol, 0.1 mM PMSF, 0.3 mg/ml bovine serum albumin, 10% glycerol, 0.25 mg/ml poly(dI-dC)/poly(dI-dC). We added 1 μl of nuclear extract and incubated samples at room temperature for 30 minutes. We loaded samples onto a 4% acrylamide (mono/bis, 29:1)-0.5× TBE-2.5% glycerol slab gel. We performed electrophoresis at 4°C, 180 V for 3-4 hours using 0.5× TBE-2.5% glycerol as a running buffer. We dried gels and imaged using a Typhoon 9410 scanner and Image Gauge software.

## Supplemental Data

**Figure S1.**
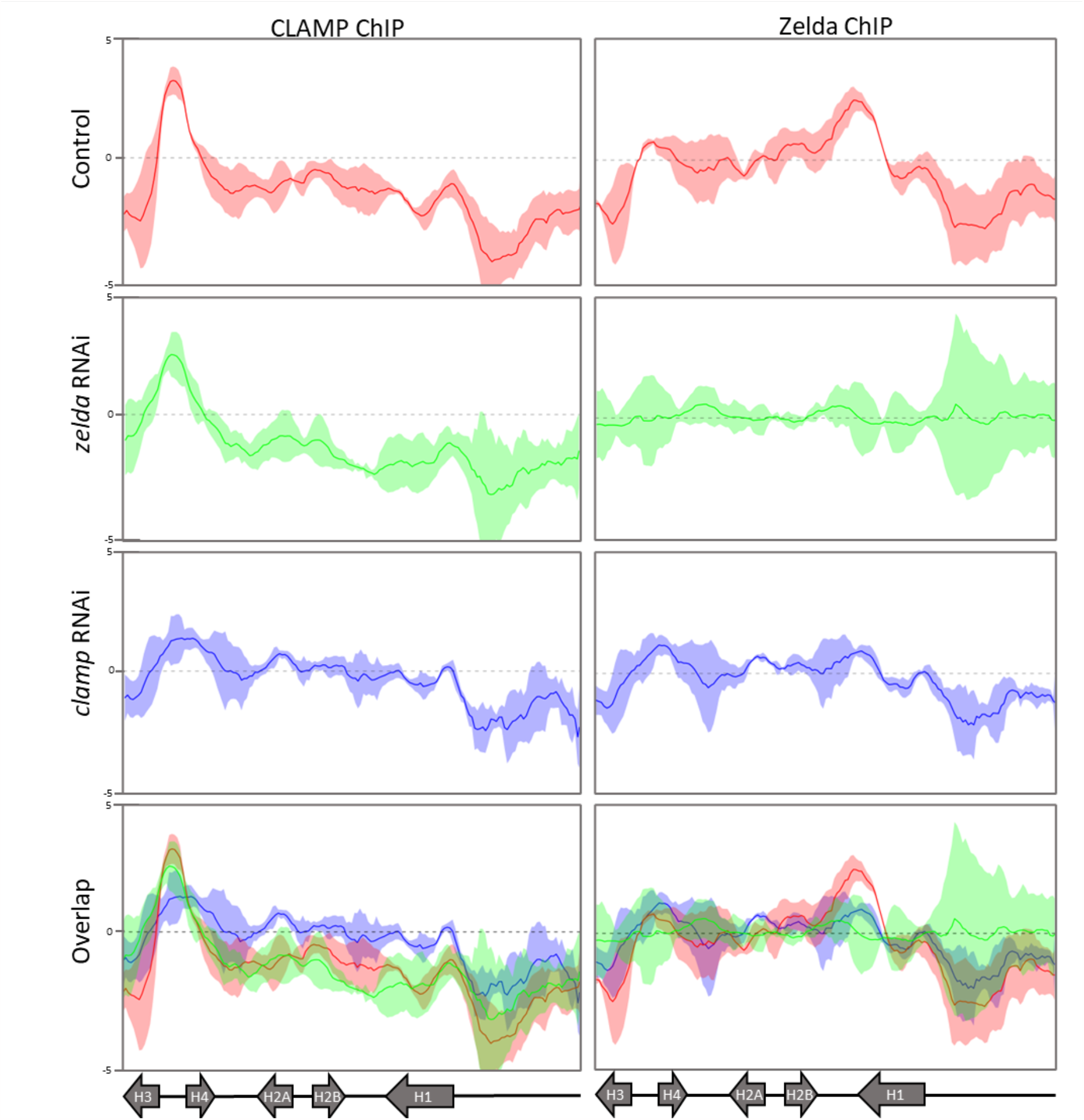
Zelda and CLAMP RNAi in the early embryo do not decrease the other’s binding at the histone locus. We mapped CLAMP and Zelda ChIP-seq data from *Drosophila* embryos 2-4hrs post lay to the histone locus in control, Zelda depleted, and CLAMP depleted conditions. CLAMP ChIP-seq shows that CLAMP localizes to the *histone3/histone4* promoter (where CLAMP’s binding motif, GA repeats, reside) and remains there under Zelda depletion and CLAMP depletion, indicating that 1) the histone locus is able to retain important factors even when they are reduced in the embryo and 2) CLAMP binding at the histone locus is not dependent on Zelda. Zelda ChIP-seq shows that Zelda binds across the locus but is mostly lost after Zelda RNAi and has slight distribution changes after CLAMP RNAi, but is mostly unaffected.

**Figure S2.**
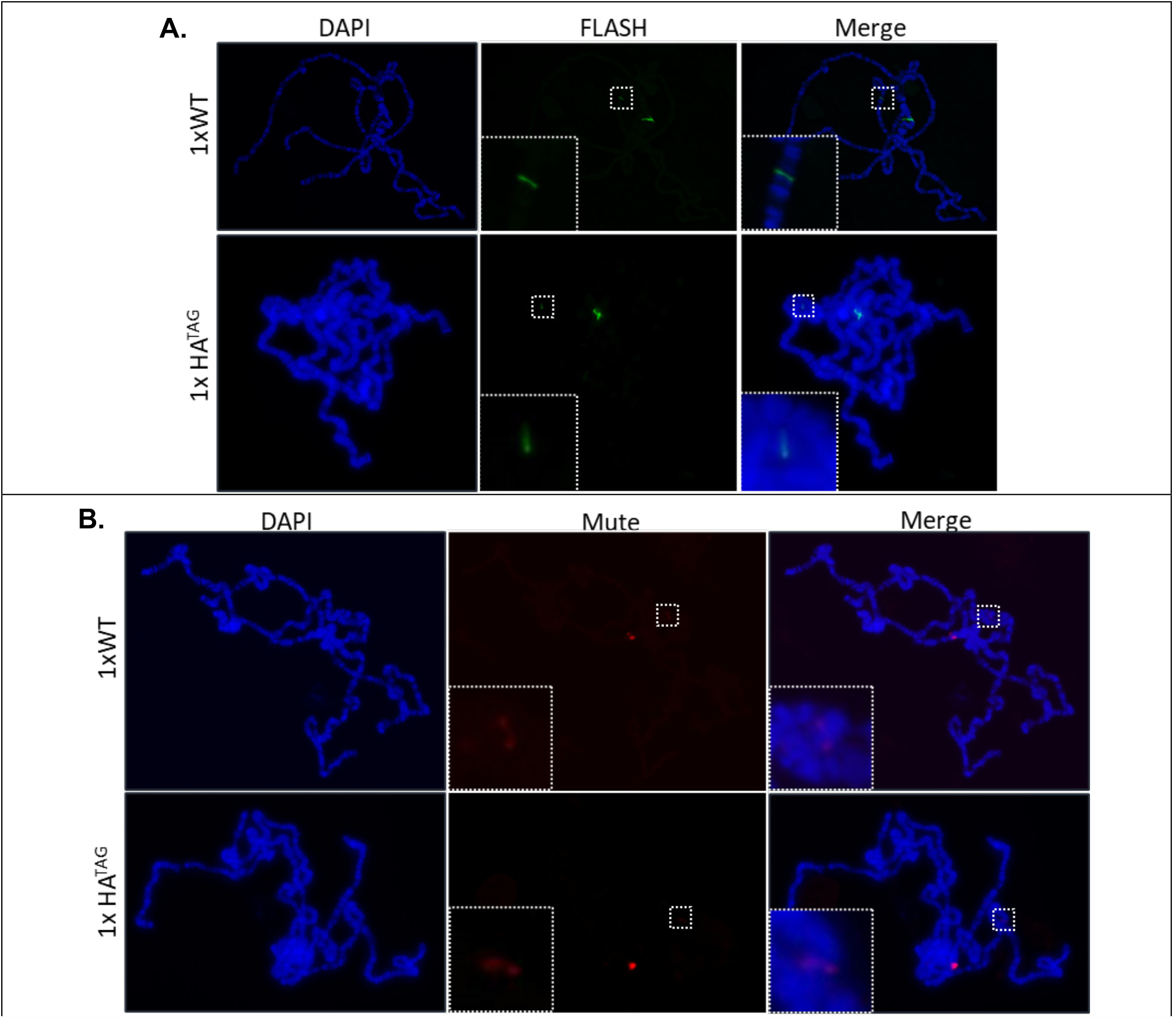
Multiple HLB factors continue to localize to transgenic histone locus arrays lacking TAGteam sites. **(A)** Polytene chromosome immunostaining for the HLB factor FLASH continues to show that histone arrays lacking some TAGteam sites (1xHA^TAG^) recruit HLB factors. **(B)** Polytene chromosome immunostaining for the HLB factor Mute.

**Figure S3.**
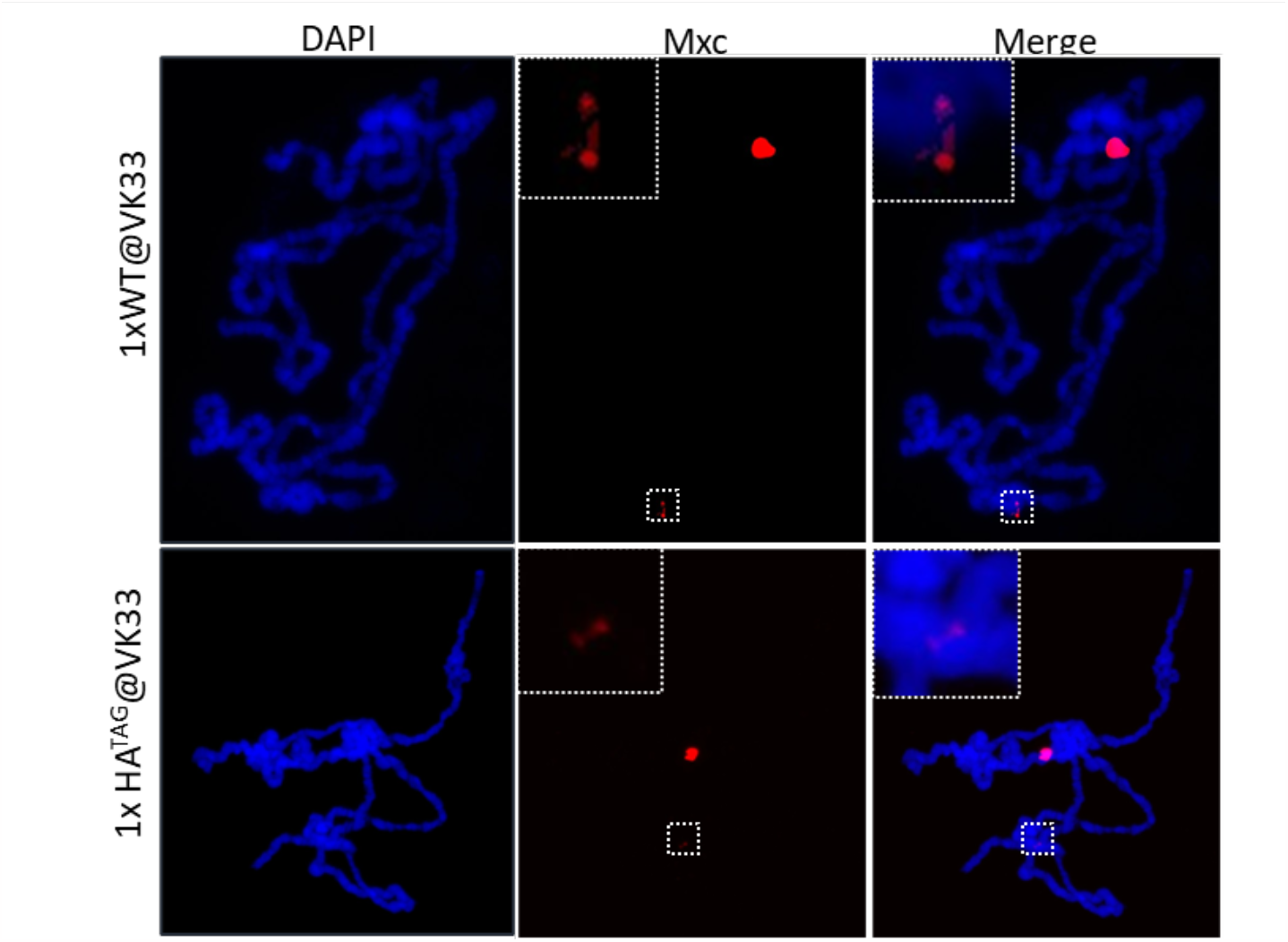
Transgenic histone loci show varying fluorescent strength at different genomic insertion sites. Polytene chromosome immunostaining on glands extracted from a fly line in which the 1xWT and 1xHA^TAG^ were inserted into the genomic site, VK33. The fluorescence from these transgenic arrays is decreased compared to bands compared to when using the insertion site Zh86-Fb. Overexposure is required to visualize ectopic bands leading to an overexposed endogenous locus.

**Figure S4.**
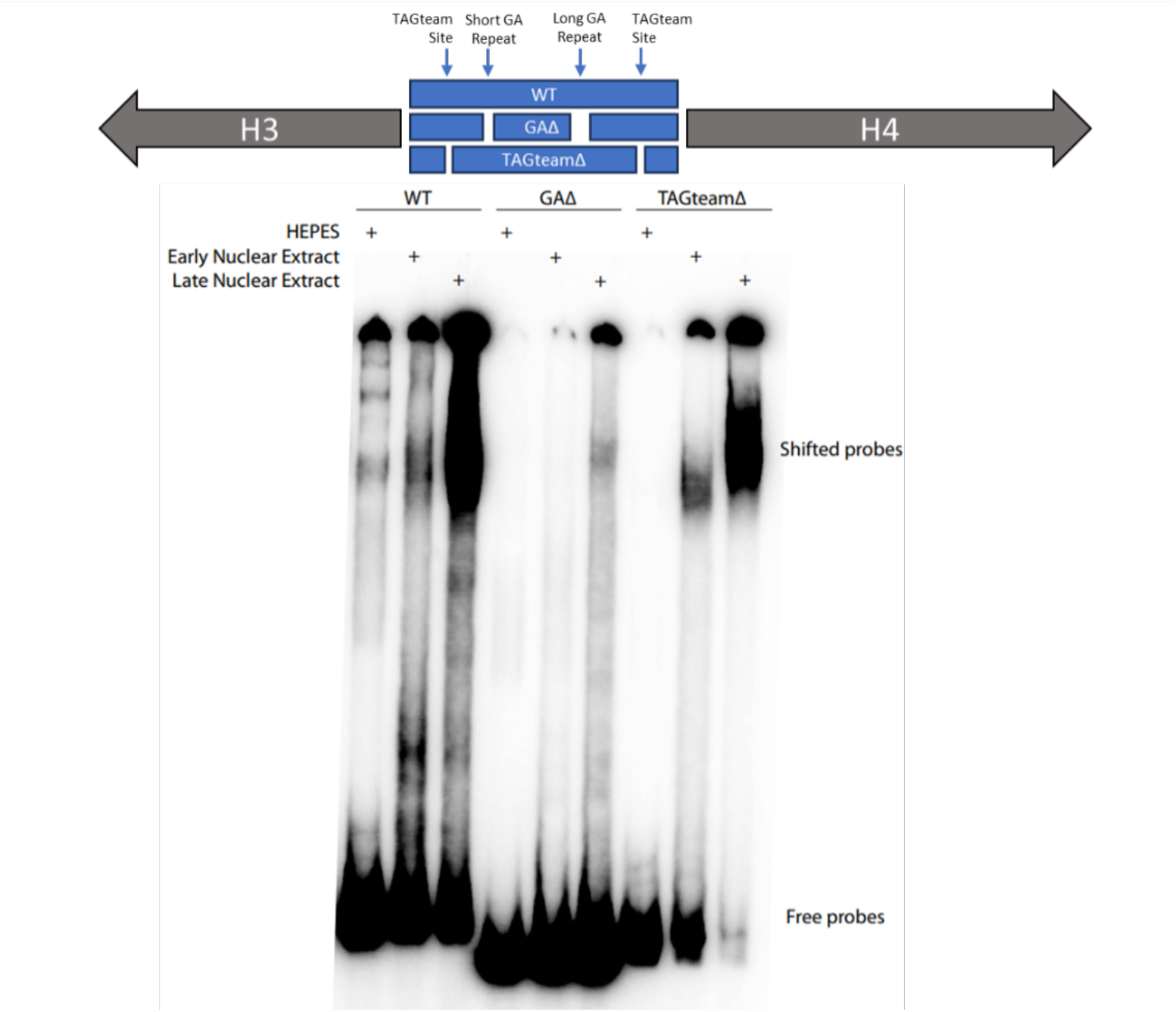
Histone array probes lacking Zelda binding sites are still bound by early embryonic proteins while probes lacking GA repeats are not. We performed an electrophoretic mobility shift assay (EMSA) using DNA probes that we designed to match regions of the *histone3/histone4* promoter. We used probes that were representative of the wild-type (WT) sequence, lacked GA repeats (GAΔ, the binding sites for CLAMP), and lacked the TAGteam sites (TAGteamΔ, the binding sites for Zelda). We exposed these probes to early (0-12hr) and late (12-24hr) embryo nuclear extract. We observe that the GAΔ probes have greatly reduced shifting compared to the WT probes indicating a loss of protein interaction. We see that the degree of shifting is mostly maintained when using the ZeldaΔ probes indicating that the binding of Zelda is not required for other HLB factors to recognize and bind the histone locus sequence.

**Table S1.**
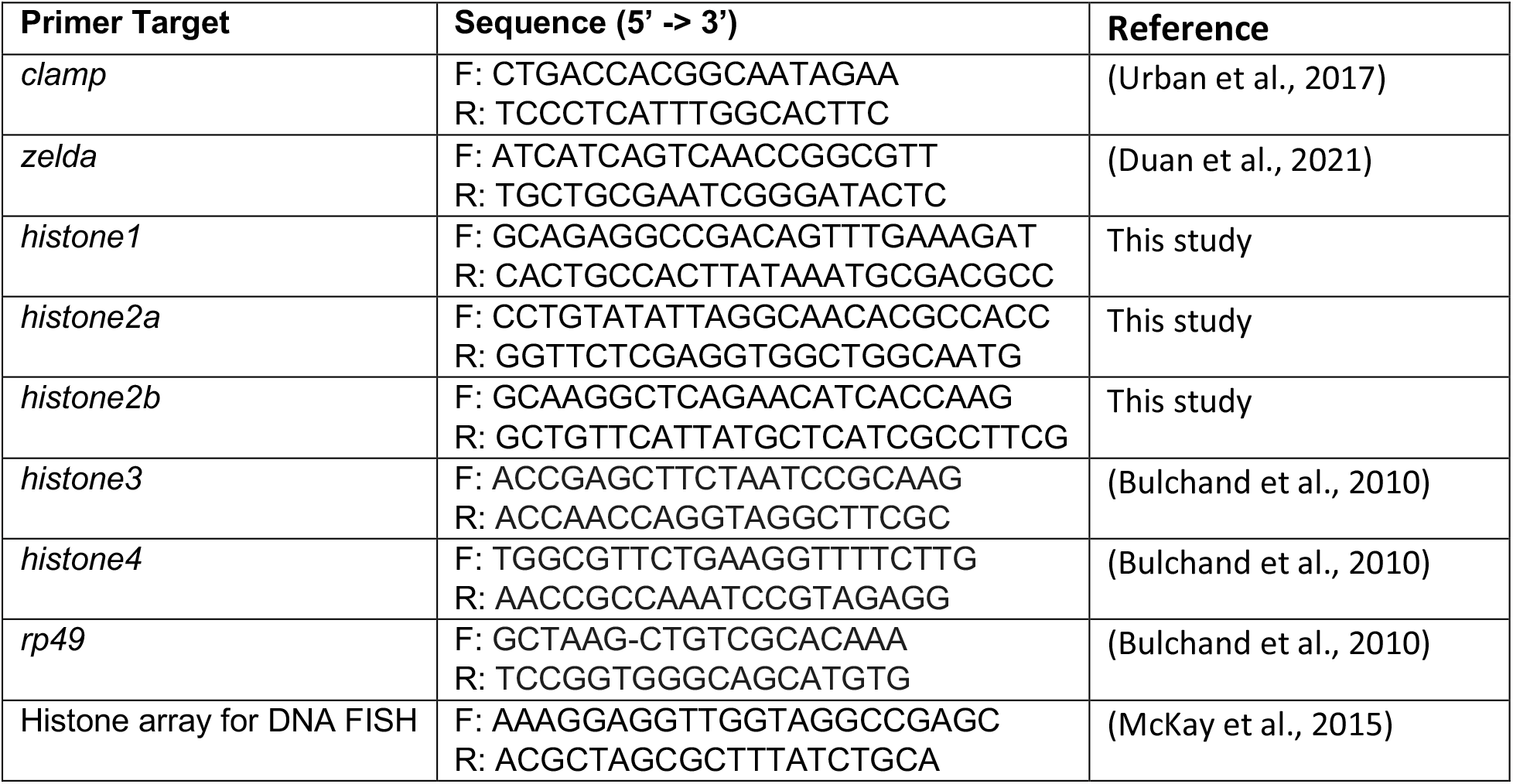

## Acknowledgments

We would like to thank Drs. Robert Duronio and William Marzluff for the anti-Mcx antibody, Dr. Chris Rushlow for the anti-*zelda* shRNA line, and Dr. Melissa Harrison for the Zelda-sfGFP stock. We thank laboratory members for their critical reading of the manuscript.

## References

Afgan, E., Baker, D., van den Beek, M., Blankenberg, D., Bouvier, D., Čech, M., Chilton, J., Clements, D., Coraor, N., Eberhard, C., Grüning, B., Guerler, A., Hillman-Jackson, J., Von Kuster, G., Rasche, E., Soranzo, N., Turaga, N., Taylor, J., Nekrutenko, A., & Goecks, J. (2016). The Galaxy platform for accessible, reproducible and collaborative biomedical analyses: 2016 update. Nucleic Acids Res, 44(W1), W3–w10. 10.1093/nar/gkw343

Amodeo, A. A., Jukam, D., Straight, A. F., & Skotheim, J. M. (2015). Histone titration against the genome sets the DNA-to-cytoplasm threshold for the Xenopus midblastula transition. Proceedings of the National Academy of Sciences of the United States of America, 112(10), E1086–E1095. 10.1073/PNAS.1413990112/-/DCSUPPLEMENTAL

Annunziato, A. (2008). DNA packaging: nucleosomes and chromatin. Nature education, 1(1), 26.

Aoki, T., Schweinsberg, S., Manasson, J., & Schedl, P. (2008). A stage-specific factor confers Fab-7 boundary activity during early embryogenesis in Drosophila. Mol Cell Biol, 28(3), 1047–1060. 10.1128/mcb.01622-07

Bongartz, P., & Schloissnig, S. (2019). Deep repeat resolution—the assembly of the Drosophila Histone Complex. Nucleic Acids Research, 47(3), e18–e18. 10.1093/NAR/GKY1194

Bulchand, S., Menon, S. D., George, S. E., & Chia, W. (2010). Muscle wasted: A novel component of the Drosophila histone locus body required for muscle integrity. Journal of Cell Science, 123(16), 2697–2707. 10.1242/jcs.063172

Chari, S., Wilky, H., Govindan, J., & Amodeo, A. A. (2019). Histone concentration regulates the cell cycle and transcription in early development. Development (Cambridge), 146(19). 10.1242/dev.177402

Cho, C. Y., Kemp, J. P., Jr., Duronio, R. J., & O’Farrell, P. H. (2022). Coordinating transcription and replication to mitigate their conflicts in early Drosophila embryos. Cell Rep, 41(3), 111507. 10.1016/j.celrep.2022.111507

De Renzis, S., Elemento, O., Tavazoie, S., & Wieschaus, E. F. (2007). Unmasking activation of the zygotic genome using chromosomal deletions in the Drosophila embryo. PLoS Biol, 5(5), e117. 10.1371/journal.pbio.0050117

Duan, J. E., Rieder, L. E., Colonnetta, M. M., Huang, A., McKenney, M., Watters, S., Deshpande, G., Jordan, W. T., Fawzi, N. L., & Larschan, E. N. (2021). Clamp and zelda function together to promote drosophila zygotic genome activation. eLife, 10. 10.7554/ELIFE.69937

Dufourt, J., Trullo, A., Hunter, J., Fernandez, C., Lazaro, J., Dejean, M., Morales, L., Nait-Amer, S., Schulz, K. N., Harrison, M. M., Favard, C., Radulescu, O., & Lagha, M. (2018). Temporal control of gene expression by the pioneer factor Zelda through transient interactions in hubs. Nature Communications, 9(1). 10.1038/s41467-018-07613-z

Duronio, R. J., & Marzluff, W. F. (2017). Coordinating cell cycle-regulated histone gene expression through assembly and function of the Histone Locus Body. In RNA Biology (Vol. 14, pp. 726-738): Taylor and Francis Inc.

Edgar, B. A., & Schubiger, G. (1986). Parameters controlling transcriptional activation during early Drosophila development. Cell, 44(6), 871–877. 10.1016/0092-8674(86)90009-7

Foo, S. M., Sun, Y., Lim, B., Ziukaite, R., O’Brien, K., Nien, C. Y., Kirov, N., Shvartsman, S. Y., & Rushlow, C. A. (2014). Zelda potentiates morphogen activity by increasing chromatin accessibility. Curr Biol, 24(12), 1341–1346. 10.1016/j.cub.2014.04.032

Günesdogan, U., Jäckle, H., & Herzig, A. (2010). A genetic system to assess in vivo the functions of histones and histone modifications in higher eukaryotes. EMBO Reports, 11(10), 772–776. 10.1038/embor.2010.124

Hamm, D. C., & Harrison, M. M. (2018). Regulatory principles governing the maternal-to-zygotic transition: Insights from Drosophila melanogaster. In Open Biology (Vol. 8): Royal Society Publishing.

Hamm, D. C., Larson, E. D., Nevil, M., Marshall, K. E., Bondra, E. R., & Harrison, M. M. (2017). A conserved maternal-specific repressive domain in Zelda revealed by Cas9-mediated mutagenesis in Drosophila melanogaster. PLoS Genet, 13(12), e1007120. 10.1371/journal.pgen.1007120

Hamm, D. C., Larson, E. D., Nevil, M., Marshall, K. E., Bondra, E. R., & Harrison, M. M. (2017). A conserved maternal-specific repressive domain in Zelda revealed by Cas9-mediated mutagenesis in Drosophila melanogaster. PLOS Genetics, 13(12). 10.1371/journal.pgen.1007120

Harrison, M. M., Li, X. Y., Kaplan, T., Botchan, M. R., & Eisen, M. B. (2011). Zelda binding in the early Drosophila melanogaster embryo marks regions subsequently activated at the maternal-to-zygotic transition. PLOS Genetics, 7(10). 10.1371/journal.pgen.1002266

Hodkinson, L. J., Smith, C., Comstra, H. S., Albanese, E. H., Ajani, B. A., Arsalan, K., Daisson, A. P., Forrest, K. B., Fox, E. H., Guerette, M. R., Khan, S., Koenig, M. P., Lam, S., Lewandowski, A. S., Mahoney, L. J., Manai, N., Miglay, J., Miller, B. A., Milloway, O., … Rieder, L. E. (2023). A bioinformatics screen reveals Hox and chromatin remodeling factors at the Drosophila histone locus. bioRxiv. 10.1101/2023.01.06.523008

Horard, B., & Loppin, B. (2015). Histone storage and deposition in the early Drosophila embryo. Chromosoma, 124(2), 163–175. 10.1007/s00412-014-0504-7

Huang, S. K., Whitney, P. H., Dutta, S., Shvartsman, S. Y., & Rushlow, C. A. (2021). Spatial organization of transcribing loci during early genome activation in Drosophila. Current Biology, 31(22), 5102-5110.e5105. 10.1016/J.CUB.2021.09.027

Hug, C. B., Grimaldi, A. G., Kruse, K., & Vaquerizas, J. M. (2017). Chromatin Architecture Emerges during Zygotic Genome Activation Independent of Transcription. Cell, 169, 216–228. 10.1016/j.cell.2017.03.024

Kaye, E. G., Kurbidaeva, A., Wolle, D., Aoki, T., Schedl, P., & Larschan, E. (2017). Drosophila Dosage Compensation Loci Associate with a Boundary-Forming Insulator Complex. Mol Cell Biol, 37(21). 10.1128/mcb.00253-17

Kemp, J. P., Yang, X. C., Dominski, Z., Marzluff, W. F., & Duronio, R. J. (2021). Superresolution light microscopy of the Drosophila histone locus body reveals a core-shell organization associated with expression of replication-dependent histone genes. Molecular Biology of the Cell, 32(9), 942–955. 10.1091/MBC.E20-10-0645

Koreski, K. P., Rieder, L. E., McLain, L. M., Chaubal, A., Marzluff, W. F., & Duronio, R. J. (2020). Drosophila histone locus body assembly and function involves multiple interactions. Molecular Biology of the Cell, 31(14), 1525–1537. 10.1091/mbc.E20-03-0176

Langmead, B., & Salzberg, S. L. (2012). Fast gapped-read alignment with Bowtie 2. Nature Methods, 9(4), 357–359. 10.1038/nmeth.1923

Larschan, E., Soruco, M. M., Lee, O. K., Peng, S., Bishop, E., Chery, J., Goebel, K., Feng, J., Park, P. J., & Kuroda, M. I. (2012). Identification of chromatin-associated regulators of MSL complex targeting in Drosophila dosage compensation. PLoS Genet, 8(7), e1002830. 10.1371/journal.pgen.1002830

Larson, E. D., Komori, H., Fitzpatrick, Z. A., Krabbenhoft, S. D., Lee, C.-Y., & Harrison, M. (2022). Premature translation of the zygotic genome activator Zelda is not sufficient to precociously activate gene expression. bioRxiv, 2022.2003.2022.485419-482022.485403.485422.485419. 10.1101/2022.03.22.485419

Li, X.-Y., & Eisen, M. (2018). Effects of the maternal factor Zelda on zygotic enhancer activity in the Drosophila embryo. In: bioRxiv.

Li, Z., Thiel, K., Thul, P. J., Beller, M., Kühnlein, R. P., & Welte, M. A. (2012). Lipid droplets control the maternal histone supply of Drosophila embryos. Curr Biol, 22(22), 2104–2113. 10.1016/j.cub.2012.09.018

Liang, H. L., Nien, C. Y., Liu, H. Y., Metzstein, M. M., Kirov, N., & Rushlow, C. (2008). The zinc-finger protein Zelda is a key activator of the early zygotic genome in Drosophila. Nature, 456(7220), 400–403. 10.1038/nature07388

Liu, J. L., Murphy, C., Buszczak, M., Clatterbuck, S., Goodman, R., & Gall, J. G. (2006). The Drosophila melanogaster Cajal body. J Cell Biol, 172(6), 875–884. 10.1083/jcb.200511038

Marzluff, W. F., Gongidi, P., Woods, K. R., Jin, J., & Maltais, L. J. (2002). The Human and Mouse Replication-Dependent Histone Genes. Genomics, 80(5), 487–498. 10.1006/geno.2002.6850

McCleland, M. L., Shermoen, A. W., & O’Farrell, P. H. (2009). DNA replication times the cell cycle and contributes to the mid-blastula transition in <i>Drosophila</i> embryos. Journal of Cell Biology, 187(1), 7–14. 10.1083/jcb.200906191

McDaniel, S. L., Gibson, T. J., Schulz, K. N., Fernandez Garcia, M., Nevil, M., Jain, S. U., Lewis, P. W., Zaret, K. S., & Harrison, M. M. (2019). Continued Activity of the Pioneer Factor Zelda Is Required to Drive Zygotic Genome Activation. Molecular Cell, 74(1), 185-195.e184. 10.1016/j.molcel.2019.01.014

McKay, D. J., Klusza, S., Penke, T. J. R., Meers, M. P., Curry, K. P., McDaniel, S. L., Malek, P. Y., Cooper, S. W., Tatomer, D. C., Lieb, J. D., Strahl, B. D., Duronio, R. J., & Matera, A. G. (2015). Interrogating the function of metazoan histones using engineered gene clusters. Developmental Cell, 32(3), 373–386. 10.1016/j.devcel.2014.12.025

Mei, Q., Huang, J., Chen, W., Tang, J., Xu, C., Yu, Q., Cheng, Y., Ma, L., Yu, X., & Li, S. (2017). Regulation of DNA replication-coupled histone gene expression. Oncotarget, 8(55), 95005–95022. 10.18632/ONCOTARGET.21887

Ni, J. Q., Zhou, R., Czech, B., Liu, L. P., Holderbaum, L., Yang-Zhou, D., Shim, H. S., Tao, R., Handler, D., Karpowicz, P., Binari, R., Booker, M., Brennecke, J., Perkins, L. A., Hannon, G. J., & Perrimon, N. (2011). A genome-scale shRNA resource for transgenic RNAi in Drosophila. Nature Methods, 8(5), 405–407. 10.1038/nmeth.1592

Nien, C.-Y., Liang, H.-L., Butcher, S., Sun, Y., Fu, S., Gocha, T., Kirov, N., Manak, J. R., & Rushlow, C. (2011). Temporal Coordination of Gene Networks by Zelda in the Early Drosophila Embryo. PLOS Genetics, 7(10), e1002339–e1002339. 10.1371/journal.pgen.1002339

Rao, X., Huang, X., Zhou, Z., & Lin, X. (2013). An improvement of the 2^(-delta delta CT) method for quantitative real-time polymerase chain reaction data analysis. Biostat Bioinforma Biomath, 3(3), 71–85.

Rieder, L. E., Koreski, K. P., Boltz, K. A., Kuzu, G., Urban, J. A., Bowman, S. K., Zeidman, A., Jordan, W. T., Tolstorukov, M. Y., Marzluff, W. F., Duronio, R. J., & Larschan, E. N. (2017). Histone locus regulation by the Drosophila dosage compensation adaptor protein CLAMP. Genes and Development, 31(14), 1494–1508. 10.1101/gad.300855.117

Roberts, S. B., Sanicola, M., Emmons, S. W., & Childs, G. (1987). Molecular characterization of the histone gene family of Caenorhabditis elegans. J Mol Biol, 196(1), 27–38. 10.1016/0022-2836(87)90508-0

Robinson, J. T., Thorvaldsdóttir, H., Winckler, W., Guttman, M., Lander, E. S., Getz, G., & Mesirov, J. P. (2011). Integrative genomics viewer. Nat Biotechnol, 29(1), 24–26. 10.1038/nbt.1754

Romeo, V., & Schümperli, D. (2016). Cycling in the nucleus: regulation of RNA 3’ processing and nuclear organization of replication-dependent histone genes. Current Opinion in Cell Biology, 40, 23–31. 10.1016/J.CEB.2016.01.015

Salzler, H. R., Tatomer, D. C., Malek, P. Y., McDaniel, S. L., Orlando, A. N., Marzluff, W. F., & Duronio, R. J. (2013). A Sequence in the Drosophila H3-H4 Promoter Triggers Histone Locus Body Assembly and Biosynthesis of Replication-Coupled Histone mRNAs. Developmental Cell, 24(6), 623–634. 10.1016/j.devcel.2013.02.014

Satija, R., & Bradley, R. K. (2012). The TAGteam motif facilitates binding of 21 sequencespecific transcription factors in the Drosophila embryo. Genome Research, 22(4), 656–665. 10.1101/gr.130682.111

Satija, R., & Bradley, R. K. (2012). The TAGteam motif facilitates binding of 21 sequence-specific transcription factors in the Drosophila embryo. Genome Res, 22(4), 656–665. 10.1101/gr.130682.111

Schulz, K. N., & Harrison, M. M. (2019). Mechanisms regulating zygotic genome activation. In Nature Reviews Genetics (Vol. 20, pp. 221–234): Nature Publishing Group.

Seal, R. L., Denny, P., Bruford, E. A., Gribkova, A. K., Landsman, D., Marzluff, W. F., McAndrews, M., Panchenko, A. R., Shaytan, A. K., & Talbert, P. B. (2022). A standardized nomenclature for mammalian histone genes. Epigenetics & Chromatin, 15(1), 34. 10.1186/s13072-022-00467-2

Sun, Y., Nien, C. Y., Chen, K., Liu, H. Y., Johnston, J., Zeitlinger, J., & Rushlow, C. (2015). Zelda overcomes the high intrinsic nucleosome barrier at enhancers during Drosophila zygotic genome activation. Genome Research, 25(11), 1703–1714. 10.1101/gr.192542.115

Tadros, W., & Lipshitz, H. D. (2009). The maternal-to-zygotic transition: A play in two acts. Development, 136(18), 3033–3042. 10.1242/dev.033183

Tatomer, D. C., Terzo, E., Curry, K. P., Salzler, H., Sabath, I., Zapotoczny, G., McKay, D. J., Dominski, Z., Marzluff, W. F., & Duronio, R. J. (2016). Concentrating pre-mRNA processing factors in the histone locus body facilitates efficient histone mRNA biogenesis. Journal of Cell Biology, 213(5), 557–570. 10.1083/jcb.201504043

Terzo, E. A., Lyons, S. M., Poulton, J. S., Temple, B. R. S., Marzluff, W. F., & Duronio, R. J. (2015). Distinct self-interaction domains promote Multi Sex Combs accumulation in and formation of the Drosophila histone locus body. Molecular Biology of the Cell, 26(8), 1559–1574. 10.1091/mbc.E14-10-1445

Urban, J. A., Doherty, C. A., Jordan, W. T., 3rd, Bliss, J. E., Feng, J., Soruco, M. M., Rieder, L. E., Tsiarli, M. A., & Larschan, E. N. (2017). The essential Drosophila CLAMP protein differentially regulates non-coding roX RNAs in male and females. Chromosome Res, 25(2), 101–113. 10.1007/s10577-016-9541-9

White, A. E., Burch, B. D., Yang, X. C., Gasdaska, P. Y., Dominski, Z., Marzluff, W. F., & Duronio, R. J. (2011). Drosophila histone locus bodies form by hierarchical recruitment of components. Journal of Cell Biology, 193(4), 677–694. 10.1083/jcb.201012077

White, A. E., Leslie, M. E., Calvi, B. R., Marzluff, W. F., & Duronio, R. J. (2007). Developmental and cell cycle regulation of the Drosophila histone locus body. Molecular Biology of the Cell, 18(7), 2491–2502. 10.1091/mbc.E06-11-1033

Xie, M., Hodkinson, L. J., Comstra, H. S., Diaz-Saldana, P. P., Gilbonio, H. E., Gross, J. L., Chavez, R. M., Puckett, G. L., Aoki, T., Schedl, P., & Rieder, L. E. (2022). MSL2 targets histone genes in <em>Drosophila virilis</em>. bioRxiv, 2022.2012.2014.520423. 10.1101/2022.12.14.520423

Yamada, S., Whitney, P. H., Huang, S. K., Eck, E. C., Garcia, H. G., & Rushlow, C. A. (2019). The Drosophila Pioneer Factor Zelda Modulates the Nuclear Microenvironment of a Dorsal Target Enhancer to Potentiate Transcriptional Output. Current Biology, 29(8), 1387-1393.e1385. 10.1016/j.cub.2019.03.019

